# Dissecting Transition Cells from Single-cell Transcriptome Data through Multiscale Stochastic Dynamics

**DOI:** 10.1101/2021.03.07.434281

**Authors:** Peijie Zhou, Shuxiong Wang, Tiejun Li, Qing Nie

**Affiliations:** LMAM and School of Mathematical Sciences, Peking University, Beijing 100871, China; Department of Mathematics, University of California, Irvine, Irvine, CA 92697, USA; Department of Cell and Developmental Biology, University of California, Irvine, Irvine, CA 92697, USA

## Abstract

Advances of single-cell technologies allow scrutinizing of heterogeneous cell states, however, analyzing transitions from snap-shot single-cell transcriptome data remains challenging. To investigate cells with transient properties or mixed identities, we present MuTrans, a method based on multiscale reduction technique for the underlying stochastic dynamical systems that prescribes cell-fate transitions. By iteratively unifying transition dynamics across multiple scales, MuTrans constructs the cell-fate dynamical manifold that depicts progression of cell-state transition, and distinguishes meta-stable and transition cells. In addition, MuTrans quantifies the likelihood of all possible transition trajectories between cell states using the coarse-grained transition path theory. Downstream analysis identifies distinct genes that mark the transient states or drive the transitions. Mathematical analysis reveals consistency of the method with the well-established Langevin equation and transition rate theory. Applying MuTrans to datasets collected from five different single-cell experimental platforms and benchmarking with seven existing tools, we show its capability and scalability to robustly unravel complex cell fate dynamics induced by transition cells in systems such as tumor EMT, iPSC differentiation and blood cell differentiation. Overall, our method bridges data-driven and model-based approaches on cell-fate transitions at single-cell resolution.

## Introduction

Advances in single-cell transcriptome techniques allow us to inspect cell states and cell-state transitions at fine resolution (1), and the notion of *transition cells* (aka. hybrid state, or intermediate state cells) starts to draw increasing attention (2–4). Transition cells are characterized by their transient dynamics during cell-fate switch (3), or their mixed identities from multiple cell states (5), different from the well-defined meta-stable cells (6, 7) that usually express marker genes with distinct biological functions. Transition cells are conceived vital in many important biological processes, such as tissue development, blood cell generation, cancer metastasis or drug resistance (8).

Despite the rapid algorithmic progress in single-cell data analysis (9), it remains challenging to probe transition cells accurately and robustly from single-cell transcriptome datasets. Often, the transition cells are rare and dynamic, and herein difficult to be captured by static dimension-reduction methods (10). High-accuracy clustering methods (e.g. SC3 (11) and SIMLR (12)) tend to enforce distinct cell states, placing transient cells into different clusters, therefore only applicable to the cases of sharp cell-state transition (**Figure 1a, top**). While popular pseudotime ordering methods (13), such as DPT (7), Slingshot (14) and Monocle (15), presumes either discrete (**Figure 1a, top**) or continuous cell-state transition (**Figure 1a, middle**), quantitative discrimination between meta-stable and transition cells is lacking (7). Recently, soft-clustering techniques provides a way to estimate the level of “mixture” of multiple cell states (16), however, the linear or static models embedded in such approach make it difficult to capture dynamical properties of cells.

**Figure 1.**
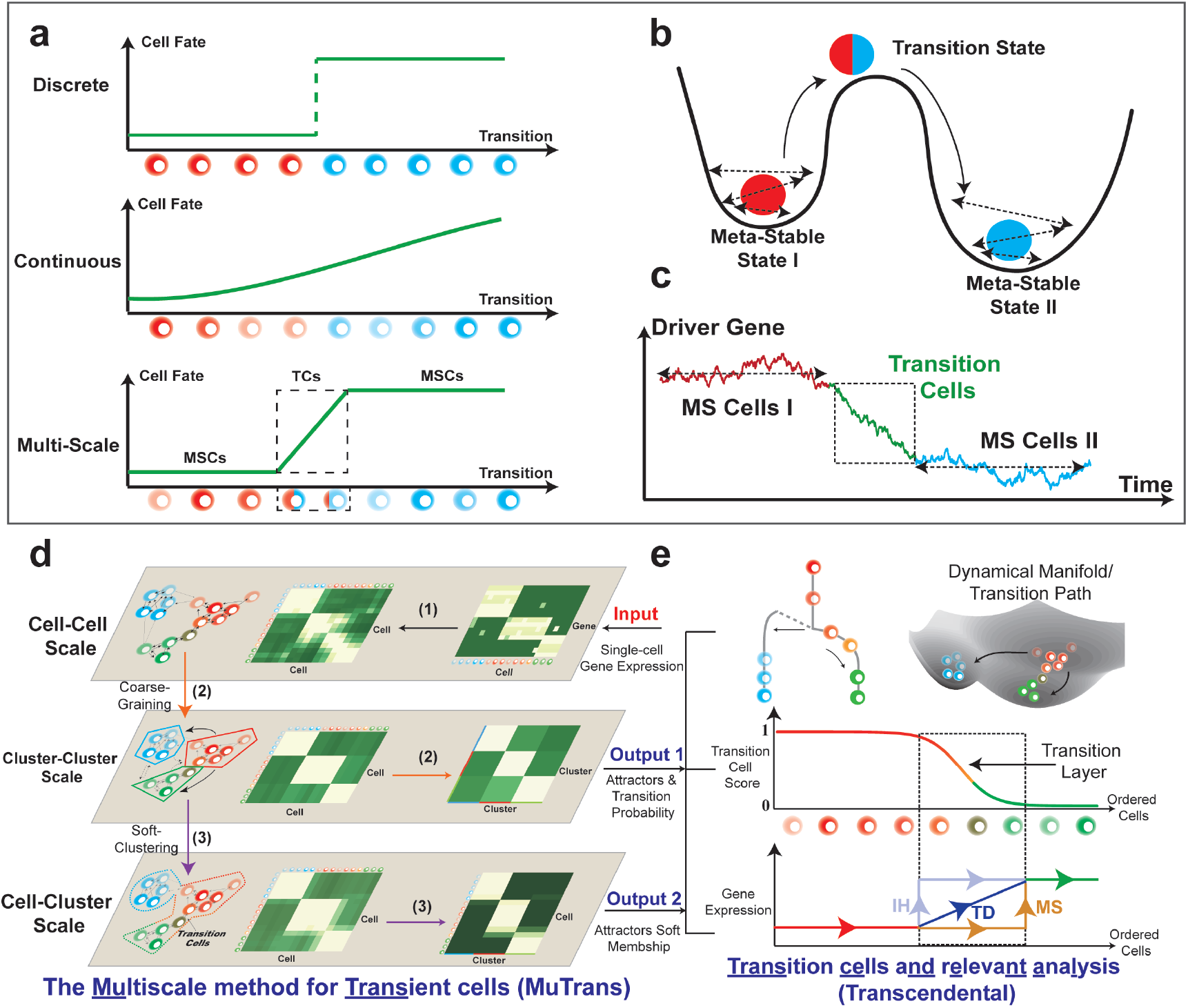
Brief introduction to MuTrans. (a-c) Theoretical foundation of MuTrans -- the multi-scale stochastic dynamics approach to model cell-fate transitions. (a) Three possible perspectives to describe cell-fate transition, as either entirely discrete (top) or continuous (middle) process, or as the multi-scale switch process between meta-stable states mediated by transition cells (bottom). The first two perspectives correspond to clustering or pseudotime ordering commonly adopted in single-cell analysis. (b) Biophysical foundation of the multi-scale perspective to treat cell-fate transition as over-damped Langevin dynamics in the multi-stable potential wells. The meta-stable states correspond to the attractor basins while the transition states are modelled by the saddle points of underlying dynamical system. (c) A typical gene expression trajectory of multi-scale dynamics. The expression of driver genes fluctuates within the meta-stable cells, while witnesses the continuous yet temporary change within transition cells, forming a transition layer in trajectory. (d-e) The procedure and downstream analysis of MuTrans. (d) The procedure of iterative multi-scale learning. The input is the preprocessed single-cell gene expression matrix. The three major steps (indicated by the number on arrow) for iterative learning of the stochastic dynamics across three different scales: (1) learning the cell-cell scale random walk transition probability matrix (rwTPM) from expression data, (2) learning the cluster-cluster scale rwTPM by coarse-graining the cell-cell scale rwTPM, and (3) learning the cell-cluster scale rwTPM by soft-clustering the cluster-cluster scale rwTPM. The output of iterative multi-scale learning includes the cell attractor basins and their mutual transition probabilities, as well as the membership matrix indicating relative cell positions in different attractors. (e) Downstream analysis (Transcendental Procedure). Given the output of iterative multi-scale learning, MuTrans constructs the cell lineage, dynamical manifold and transition paths manifesting the underlying transition dynamics of cell-fate (top). For each state-transition process, MuTrans explicitly distinguishes between meta-stable and transition cells via TCS (middle). The transition cells are marked with dashed squares. Based on the TCS ordering of cells, MuTrans identifies three types of genes (**MS, IH** and **TD**) during the transition whose expression dynamics differ in meta-stable and transition cells (bottom).

Dynamic modeling provides a natural way to characterize transition cells (3), allowing multiscale description of cell-fate transition (**Figure 1a, bottom and S1**). Such models analogize cells undergoing transition to particles confined in multiple potential wells with randomness (17, 18), for which the transient states correspond to saddle points and the metastable states correspond to attractor basins of the underlying dynamical system (**Figure 1b**). In such description, the stochastic gene dynamics at individual cell scale can induce cell-state switch at macroscopic cell cluster or phenotype scale, and the transition cells form “bridges” between meta-stable states (**Figure 1c**). Despite widely use of dynamical systems concepts to illustrate cell-fate decision (4), direct inference via dynamical models for transitions from single-cell transcriptome data is lacking.

Here we employ noise-perturbed dynamical systems (19) with a multiscale approach on cell-fate conversion (20) to analyze single-cell transcriptome data. By characterizing meta-stable cells in attractor basins and placing the transition cells along transition paths connecting the meta-stable states through saddle points, our **<u>mu</u>**ltiscale method for **<u>trans</u>**ient cells (MuTrans) prescribes a stochastic dynamical system for a given dataset (**Figure 1b**). Using the single-cell expression matrix as input, through iteratively constructing and integrating cellular random walks across three scales (**Figure 1d** and **S2**), MuTrans finds most probable path tree (MPPT) for cell transitions in a reconstructed cell-fate *dynamical manifold* (**Figure 1e**). Such manifold, similar to the classical Waddington landscape (21) often used to highlight transitions, provides an intuitive visualization of cell dynamics compared to commonly adopted low-dimension *geometrical manifold*. In the dynamical manifold, the barrier height naturally quantifies the likelihood of cell-fate switch, and a Transition Cell Score (TCS) allows us to distinguish meta-stable and transition cells (**Figure 1e**). We then illustrate the complex cell transition trajectories on dynamical manifold using the dominant transition paths obtained for the coarse-grained dynamics. With such quantification, we are able to identify critical genes that are **<u>t</u>**ransition **<u>d</u>**rivers (TD genes), mark the **<u>i</u>**ntermediate/**<u>h</u>**ybrid states (IH genes) or **<u>m</u>**eta-**<u>s</u>**table cells (MS genes) (**Figure 1e** and **S3**). To speed up calculations for datasets consisting of large number of cells (22, 23), MuTrans provides an additional (and optional) aggregation module in pre-processing. This module aggregates cells into many small groups that share similar dynamical properties, thus MuTrans can take the transition probabilities among these coarse-grained “cells” as the input, instead of the random walk on original cells, in order to reduce the computational cost (**Method and SM Section 2.6**).

We demonstrate the effectiveness and robustness of MuTrans in seven single-cell transcriptome datasets, including simulation data and sequencing data generated by five different experimental platforms. Benchmarking and comparisons with seven existing single-cell lineage inference tools validates the capability and scalability of MuTrans in probing complex, sometimes subtle, cell-fate transition dynamics. We also perform mathematical analysis to show consistency of MuTrans with the over-damped Langevin dynamics (24) -- a popular model for state transitions in physical or biochemical systems (19).

## Results

### Overview of MuTrans

MuTrans depicts cells and their transitions in a given single-cell transcriptome dataset as a multiscale dynamical system (**Figure 1a-c**). Taking the input as pre-processed single-cell gene expression matrix, MuTrans first learns the cellular random walk transition probability matrix (rwTPM) on the cell-cell scale through the Gaussian-like kernel (**Figure 1d and Methods**), which yields the continuous limit of over-damped Langevin Equation to model cell-fate decision (**Methods and Section 1 in SM**). Next, the method performs coarse-graining on the cell-cell scale rwTPM to learn the dynamics on the cluster-cluster scale, and acquires attractor basins and their mutual conversion probabilities simultaneously (**Figure 1d and Methods**). Theoretically, this step is asymptotically consistent with the Kramers’ law of reaction rate for over-damped Langevin systems (**Methods and Section 1.2 in SM**). Finally, we specify the relative position of each cell in the attractor basins with the cell-cluster resolution view of Langevin dynamics, which is constructed via optimizing a cell-cluster membership matrix (**Figure 1d and Methods**).

In the downstream analysis (Transcendental Procedure, **Figure 1e**), we construct the most probable path tree (MPPT) to infer cell lineage based on the coarse-grained transition probabilities (**Figure 1e and SM Section 2.4**). To robustly depict the lineage relationships, we use the transition path theory to quantify the likelihood of all possible transition trajectories between cell states (**Methods and Section 2.4 in SM**).

Combining the optimized cell-cluster membership matrix, MuTrans fits a dynamical manifold using mixture distribution to make meta-stable cells reside in the attractor basins while assign transition cells along the transition paths connecting different basins (**Figure 1e and Methods**), which is inspired by the Gaussian mixture approximation toward the steady-state distribution of the Fokker-Planck equation associated with the over-damped Langevin dynamics (**Methods and Section 2.3 in SM**).

For each cell-state transition, we can calculate a transition cell score (TCS) ranging between one and zero to quantitatively distinguish meta-stable and transition cells (**Figure 1e and Methods**). Finally, we systematically classify three types of genes (MS, IH and TD) during the transition whose expression dynamics differ between metastable and transition cells (**Figure 1e and Methods**). Specifically, the TD genes varies accordingly with the TCS within transition cells, and the IH genes co-express in both metastable and transition cells, while MS genes express uniquely in the meta-stable states.

To deal with the large-scale datasets, in addition to common strategies such as sub-sampling cells, we provide an option to speed up calculation by introducing a pre-processing aggregation module DECLARE (<u>d</u>ynamics-pr<u>e</u>serving <u>c</u>el<u>l</u>s <u>a</u>gg<u>re</u>gation). This module assigns the original individual cells into many (e.g. hundreds or thousands) microscopic meta-stable states and computes the transition probabilities among them, and thus it can be used as an input to MuTrans instead of the cell-cell rwTPM (**Methods and Section 2.6 in SM**). Both theoretical and numerical analysis suggest that, compared to the common strategy of averaging of gene expression profiles of a small group of cells, DECLARE better preserves the structure of dynamical landscape with a good approximation to the transition paths probabilities calculated without using DECLARE (**Figure 5, Methods and Section 2.6 in SM**).

**Figure 2.**
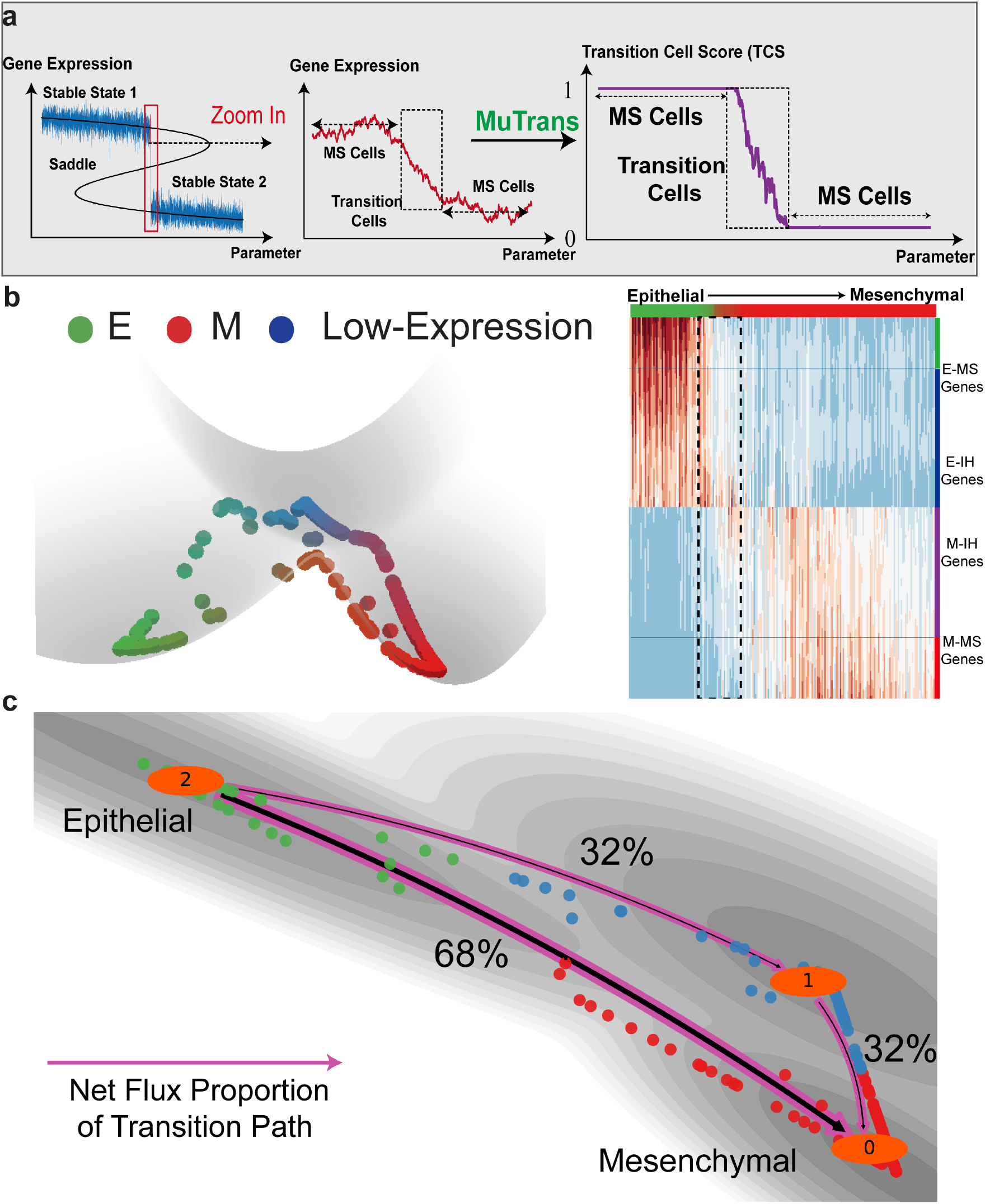
Validation of MuTrans in two-state transition simulation data and three-state EMT single-cell RNA-seq data. (a) MuTrans distinguishes the meta-stable and transition cells simulated using a stochastic saddle-bifurcation model. (top, left) The data generated by the model. (Blue lines) The simulated trajectories as the input data. (Black Lines) Bifurcation plot of the underlying dynamical system. (Red Lines) The trajectory points corresponding to the transition cells that are switching between two states. (top, right) The zoomed-in trajectory of the transition cell region. (bottom) The TCS values for transition cells. The meta-stable cells have TCS of value 0 or 1, while the TCS of transition cells decrease from 1 to 0 during transition. (b) MuTrans distinguishes between MS and IH genes, and resolves dynamics during epithelial-mesenchymal transition (EMT) mediated by transition cells. (top) The constructed dynamical manifold reveals the existence and transitions among three cell states. (bottom) The Transcendental analysis of EMT, with the genes (rows) grouped by IH or MS, is consistent with previous findings (exact names and details shown in **Table S2** and **S3**), cells (columns) ordered by TCS, and transition cells marked by the black dashed rectangles. No significant TD genes are detected during the transition. The color-map from blue to red represents low to high gene expression values. (c) The transition path analysis by setting E as start state and M as target state, overlaid on the two-dimensional dynamical manifold. The numbers are the relative likelihood of each transition path. The direct transition from E to M across the barrier of transition is the dominant path with larger transition path flux.

**Figure 3.**
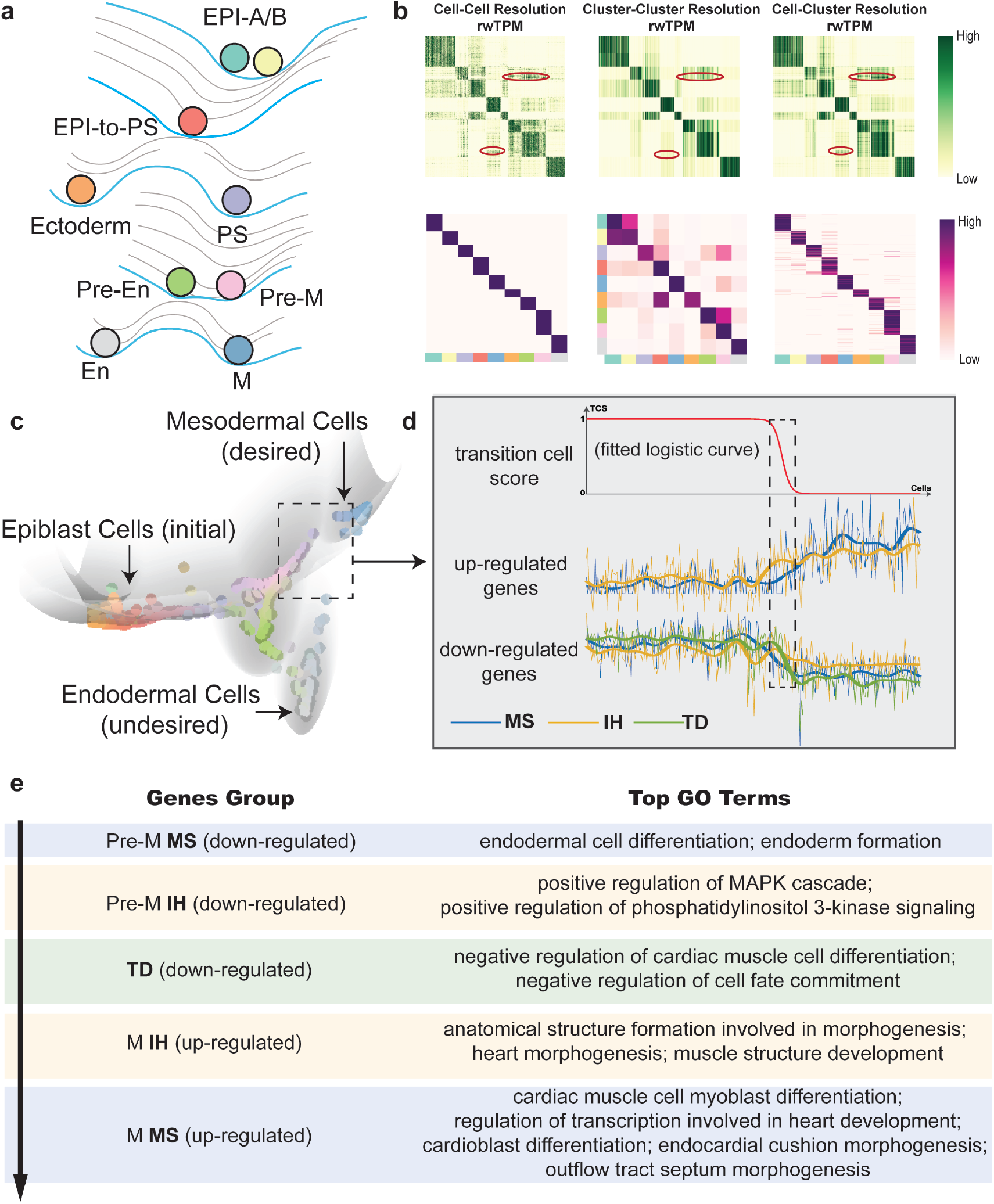
MuTrans scrutinizes the cellular bifurcation and gene expression dynamics during iPSC differentiation. (a) The schematic development landscape during iPSCs differentiation, with cell states and lineage relationship inferred by MuTrans. (b) The multi-scale quantities learned by MuTrans. (top) The learned cellular random walk transition probability matrix (rwTPM). Elements in red circle indicate that cell-cluster scale rwTPM recovers the finer resolution of cell-cell scale rwTPM than the clustercluster scale rwTPM. (bottom) The cell-cluster assignment (left), cluster-cluster transition probability (middle) and cell-cluster membership matrix (right) learned by MuTrans. (c) The constructed dynamical manifold (Methods and Section 2.3 in SM) reflects the dynamics from initial epiblast cells toward the final mesodermal (the desired cell fate in iPSC induction) or endodermal cells. The color of each individual cell is computed based on the value of its soft clustering membership. (d) The Transcendental analysis of the transition from Pre-M state to M-state (details in Section 3.3 of SM). (top) The TCS of transition, with transition cells marked by dashed rectangles. Transition cells are marked by dashed squares. (middle) The average gene expression of top 5 down-regulated MS (blue) and IH (yellow) genes. The full gene name list is shown in Table S6. The thin lines represent the raw normalized expression value and thick lines denote the smoothed data. IH genes are up-regulated in both transition and metastable M cells, while the expression of MS genes is inhibited in transition cells. (bottom) The average gene expression of top 5 down-regulated MS (blue), IH (yellow) and TD (green) genes. The full gene name list is shown in Table S6. (e) GO enrichment analysis of MS, IH and TD genes during Pre-M to M state transition indicates a gradual loss of endodermal property and gain of mesodermal property in the cell-fate switch.

**Figure 4.**
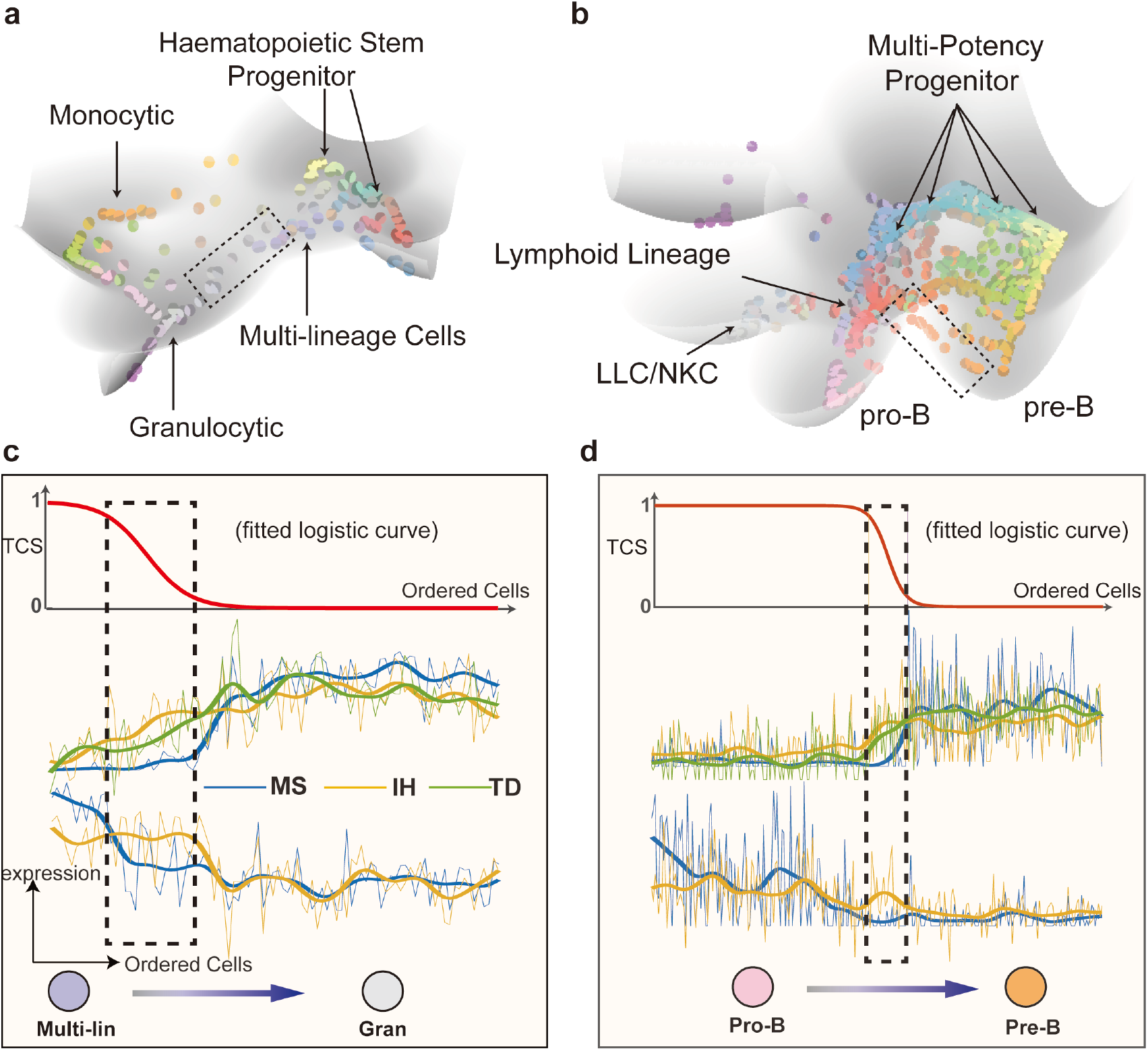
MuTrans can robustly reveal the underlying complex dynamics in single-cell blood differentiation datasets. (a-b) The constructed dynamical manifold by MuTrans are shown for the two datasets. The color of each individual cell in dynamical manifold is based on its soft-clustering membership. In mouse HPC dataset (left), MuTrans highlights the multi-lineage cells in a shallow pit on dynamical manifold. In the HPC dataset toward lymphoid lineages (right), MuTrans discovers plenty of transition cells exist between meta-stable PreB and B cell attractors (marked by dashed squares). (c) The TCS of transition and average gene expression of the top 5 TD (green), MS (blue) and IH (yellow) genes for the two interested transition paths marked with dash in (a). The full gene lists are shown in Table S7-9.

**Figure 5.**
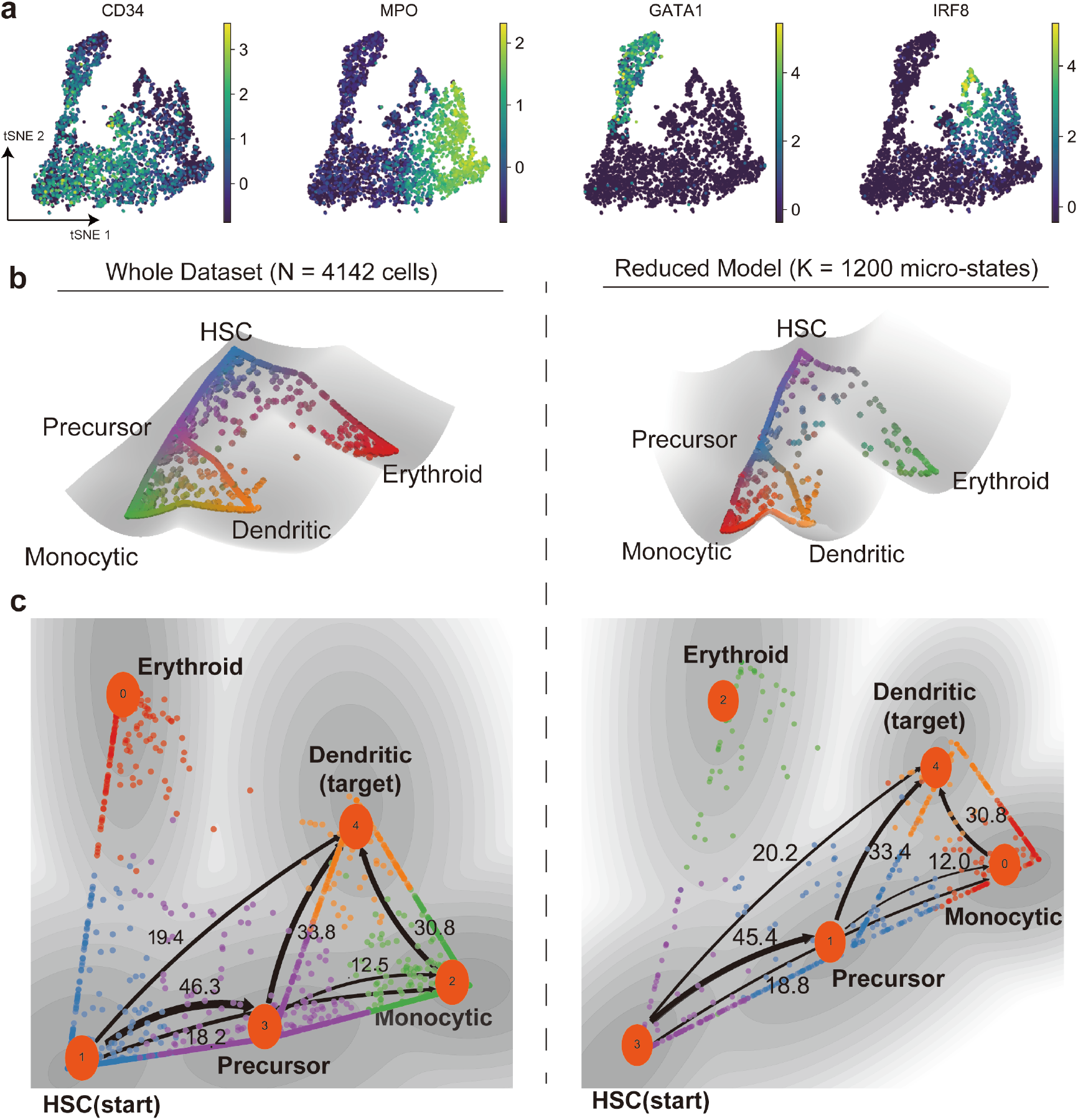
Application to a large dataset using multiscale reduction approach. (a) The tSNE plot and marker gene expression of datasets from early human HSC differentiation in bone marrow. (b) The dynamical manifold constructed from complete dataset (left, N=4,142 cells) and with DECLARE pre-processing (right, K=1,200 micro-states) with cells colored by soft clustering membership in MuTrans attractors. Left panel: each ball represents one cell; right panel: each ball represents one microstate. The reduced model preserves the overall structure of dynamical manifold. (c) The transition paths analysis conducted on complete data (left) and with DECLARE preprocessing (right), where HSC are picked as the start and dendritic cells as the target. The numbers indicate the relative likelihood of each transition path, suggesting the quantitative consistency of reduced model with the analysis on whole dataset.

### Validation in two-state simulation data and three-state EMT system

We first validated the performance of MuTrans on single-cell data generated from relatively simple cell-state transition dynamics. To test accuracy and robustness of our method, we simulated the stochastic state-transition process using a bifurcation model in the regime of intermediate noise level (25). The gene expression of each cell was simulated with over-damped Langevin equation driven by an extrinsic signal and noise (**Section 3.1 in SM**). In certain parameter range, the model consists of two stable states and one unstable saddle states (**Figure 2a**). Noise in gene expression induced the switch prior to the bifurcation point, resulting in a thin layer of transition cells (**Figure 2a**). Applying MuTrans to the known transition cells and meta-stable cells in the model, we found the computed transition cell score (TCS) captured the underlying saddle-node bifurcation structure (**Figure 2a**). For cells fluctuating around the two stable branches, the TCS approaches one or zero respectively, indicating the meta-stability of cell states. The transition cells that surpasses the saddle point region in the trajectory yields a continuum of TCS between zero and one, with scores consistent with the relative positions of cells along the trajectory (**Figure 2a**).

We then applied MuTrans to a single-cell RNA sequencing dataset (26) of tumor epithelial-to-mesenchymal transition (EMT) generated by Smart-Seq2 platform (**Figure 2b and S4-S7**). Three cell states were detected, including epithelial (E) state and mesenchymal (M) state, manifesting as the adjacent basins in the dynamical manifold, with identified EMT transition cells moving in-between (**Figure 2b, Figure S4-S6**). The transition cells were characterized by the groups of IH genes without observing significant TD genes (**Figure 2b**), agreeing well with the experimentally measured “hybrid genes” of EMT cells and the role of IH in transition (26). Compared with previous selected marker genes, we identified consistent MS markers such as Epcam, Cdh1 and Mm9, and IH markers such as Trp63 and Pdgfra (**Table S4 and S5**). It is interesting to note that the previously identified hybrid gene Krt14 was assigned into the MS group (**Table S4**), however, with low statistical significance, indicating its potential resemblance with IH genes. This agrees well with an ATAC-seq analysis (26), showing the chromatin regions of Krt14 and Krt17 in transition cells, although remained open, were actually in reduced levels. The analysis also indicates that the trajectory from epithelial state to mesenchymal state mediated by transition cells has a larger probability flux than the path surpassing another low-expression state (**Figure** 3c).

### Scrutinizing bifurcation dynamics during iPSC induction

We next used MuTrans to investigate cell fate bifurcations (**Figure 3a**) in a single-cell dataset for induced pluripotent stem cells (iPSCs) toward cardiomyocytes (27). In the learned cellular random walk across different scales, the rwTPM on cell-cluster scale recovers finer resolution of rwTPM on the cell-cell scale than the cluster-cluster scale (**Figure 3b, top**). MuTrans identified nine attractor basins (**Figure 3b, bottom left)**, and the constructed most probable path tree (**MPPT, Figure S7**) reveals a lineage with bifurcation into mesodermal (M) or endodermal (En) cell fates. Two previously unfound states, located prior to the bifurcation of primitive streak (PS) into differentiated mesodermal (M) or endodermal (En) cell fates in the MPPT, were denoted as Pre-M and Pre-En states (**Figure 3b** and **S7**). On the inferred dynamical manifold (**Figure 3c**), the cells make transitions between two states, suggesting possible dynamic conversion between the two types of precursor cells that seem to be very plastic. In comparison, the transition between mature En and M states are rare, indicating the stability of En and M cells. Along the differentiation trajectory from PS to Pre-M, the coarse-grained transition probability, quantified by the heights of barrier, shows a stronger transition capability from PS to Pre-M than from Pre-M to PS (**Figure 3b** and **S7)**. In addition, the transition from Pre-M to M was found to be sharper than the one from PS to Pre-M. The transitions from PS to Pre-En and from Pre-En to En exhibit similar behavior.

Downstream analysis on gene expression profiles indicates three transition stages from Pre-M to M (**Figure 3d)**. The initial stage was characterized by downregulation of meta-stable (MS) genes from the Pre-M state markers (enriched in the pathways of endodermal development) and upregulation of intermediate-hybrid (IH) genes (enriched in pathways of MAPK cascade and metabolic process) from the M state markers (**Table S6 in SM and Figure 3e**). This process by first losing En identity enables a conversion of Pre-M meta-stable cells toward the transition cells. The second stage of the transition marked by the gradual down-regulation of TD genes mainly involves negative regulation of cardiac muscle cell differentiation and cardiac muscle tissue development (**Table S6 in SM and Figure 3e**). The final stage completes the transition process with the down-regulation of Pre-M state IH genes, along with upregulation of MS genes (enriched in the cardiac muscle cell myoblast differentiation and outflow tract morphogenesis process) in the M state (**Table S6 in SM and Figure 3e**), making transition cells to finally convert into the mesodermal cells and establish the meta-stable cell fate. The ordering of cells based on TCS has an overall increasing trend from Day 2 to Day 3 via the time point of Day 2.5 within the transition cells, corresponding to the noticed three-stage transition (**Figure S8**). Together, the transition cells locating near the saddle points connecting Pre-M (or Pre-En) and M (or En) reflect the temporal orderings of cell-fate conversion, which are well characterized by TD and IH genes in a system consisting of one pitchfork bifurcation.

### MuTrans robustly resolves complex lineage dynamics in blood cell differentiation

The hematopoiesis has been conceived as a hierarchy of discrete binary state-transitions, while increasing evidence alternatively supports a continuous and heterogeneous view of such process (28). To investigate the complex dynamics in blood differentiation where transition cells likely play key roles, we applied MuTrans to three different single-cell datasets with different sequencing depths and sample sizes.

We first analyzed the single-cell RNA data during myelopoiesis sequenced with Fluidigm C1 platform (29). Notably MuTrans highlights the hub states in the inferred MPPT cell lineage (**Figure 4a** and **Figure S10**), capable of becoming three types of blood cells through a shallow basin resided in the highest terrain of the entire dynamical manifold (**Figure S11**). The low barriers between the multi-lineage basin and the downstream basins (granulocytic or monocytic states) suggest probable transitions from the multi-lineage state, consistent with the observed transition cells across the saddle point. Interestingly, the transition cells during Multi-lin to Gran conversion were previously identified as the multi-lineage cells in ICGS clustering (29) (**Figure S11**). Similarly, during the megakaryocytic cell differentiation, while the transition cells consist of both HSPC1 and Meg types in our analysis, they were previously identified as the hematopoietic progenitor cells by the ICGS criterion (**Figure S11**). Such discrepancy could be explained by the gene expression dynamics in gradual transition of cell states. For example, during transition from multi-lineage cells to granulocytic cells (**Figure 4c**), we observed the typical expression pattern of TD, MS and IH genes as conceptualized in **Figure 1e**. Despite the similarity between the transition cells and their departing multi-lin state as manifested in the co-expression of down-regulated IH genes (bottom panel in **Figure 4c, yellow lines**), we also detected the up-regulated IH genes (middle panel in **Figure 4c, yellow lines**), suggesting the resemblance of transition cells with their targeting gran cell state (**Table S7**). We observed a similar gene expression pattern in the transition from HSPC to Meg state (**Figure S13** and **Table S8**). For this dataset, MuTrans is able to capture the established meta-stable states, in addition to finding transition cells that were classified in some meta-stable states by a previous study (29).

Focusing on the cell-fate bias toward lymphoid lineage, MuTrans resolves the complex lineage dynamics underlying single-cell RNA data of mouse hematopoietic progenitors differentiation sequenced from Cel-Seq2 platform (30). Consistent with the major finding of FateID algorithm, the constructed dynamical manifold reveals that lymphoid progenitor (LP) cells (red balls) give rise to both B cells (pink balls) and plasmacytoid dendritic cells (pDCs) (**Figure 4b** and **S14**). The inferred MPPT and dynamical manifold also suggests that certain transition cells in the attractors of pDCs originate directly from multi-potent progenitor (MPP) cells (yellow balls, **Figure S14**). Interestingly, MuTrans resolve the details in B cell differentiation, capturing the transition cells from Pro-B toward Pre-B basins (**Figure S14** and **Table S9**). Downstream analysis validated the transition cells by the co-expressed IH genes (yellow lines, **Figure 4c right**) and the dynamically expressed TD genes (green lines, **Figure 4c right**). Overall, MuTrans provides a clear global cell-fate transition picture with marked transition cells in this dataset of highly complex lineages, in contrast to the local transition routes inferred by FateID (30).

### Application to large-scale datasets with complex trajectory

To test the scalability of MuTrans, we studied on the single-cell hematopoietic differentiation data in human bone marrow generated by 10x Chromium platform (31) (**Figure 5a**). To make the comparison, we applied MuTrans to both the complete (original) data, and the one after using the pre-processing module DECLARE. We found DECLARE could reduce the calculation time by one magnitude for this dataset.

For both cases MuTrans identified the expected bifurcations from hematopoietic stem progenitor cells (HSPC) into the monocytic precursors and erythroid cells, as well as the differentiation from precursor cells into monocytic and dendritic cells. The constructed dynamical manifold (**Figure 5bc, Figure S15**) shows a continuous stream of transition cells among different basins (such as those moving between dendritic and monocytic potential wells) suggesting the hematopoietic differentiation may be a continuous process. The transition trajectories obtained with the large-scale pre-processing step are consistent with the complete dataset analysis (**Figure 5bc**). This indicates the major transition trajectories toward dendritic cell fate not only consist of the path mediated by monocytic precursor states but also include a considerable flux of transition cells from differentiated monocytic cells. Interestingly, the existence of both meta-stable states and transition cells reconciles a previously noted discrepancy (31) caused by treating the underlying cellular transition dynamics as either a purely continuous processing (e.g. using Palantir) or a discrete process (using other clustering-based lineage inference methods such as Slingshot (14) and PAGA (32)).

Next, we analyzed another dataset containing over 15,000 cells collected during blood emergence in mouse gastrulation (33) (Figure 6a). Consistent with the PAGA (32) representation of the data (Figure 6b), the constructed dynamical manifold (Figure 6c) and derived most probable flow tree (MPFT) suggest three major transition branches from haemato-endothelial (Haem) cells into endothelial cells (EC), mesoderm cells (Mes) or erythroid cells (Ery). Specifically, the transition path analysis indicates that the endothelial cells and erythroid cells are originated through discrete trajectories from haemogenic endothelium (Figure 6e), and such trajectories are mediated by the intermediate state of blood progenitor (BP) cells (Figure 6f). These results are consistent with the experimental findings on endothelial and erythroid cells (33).

**Figure 6.**
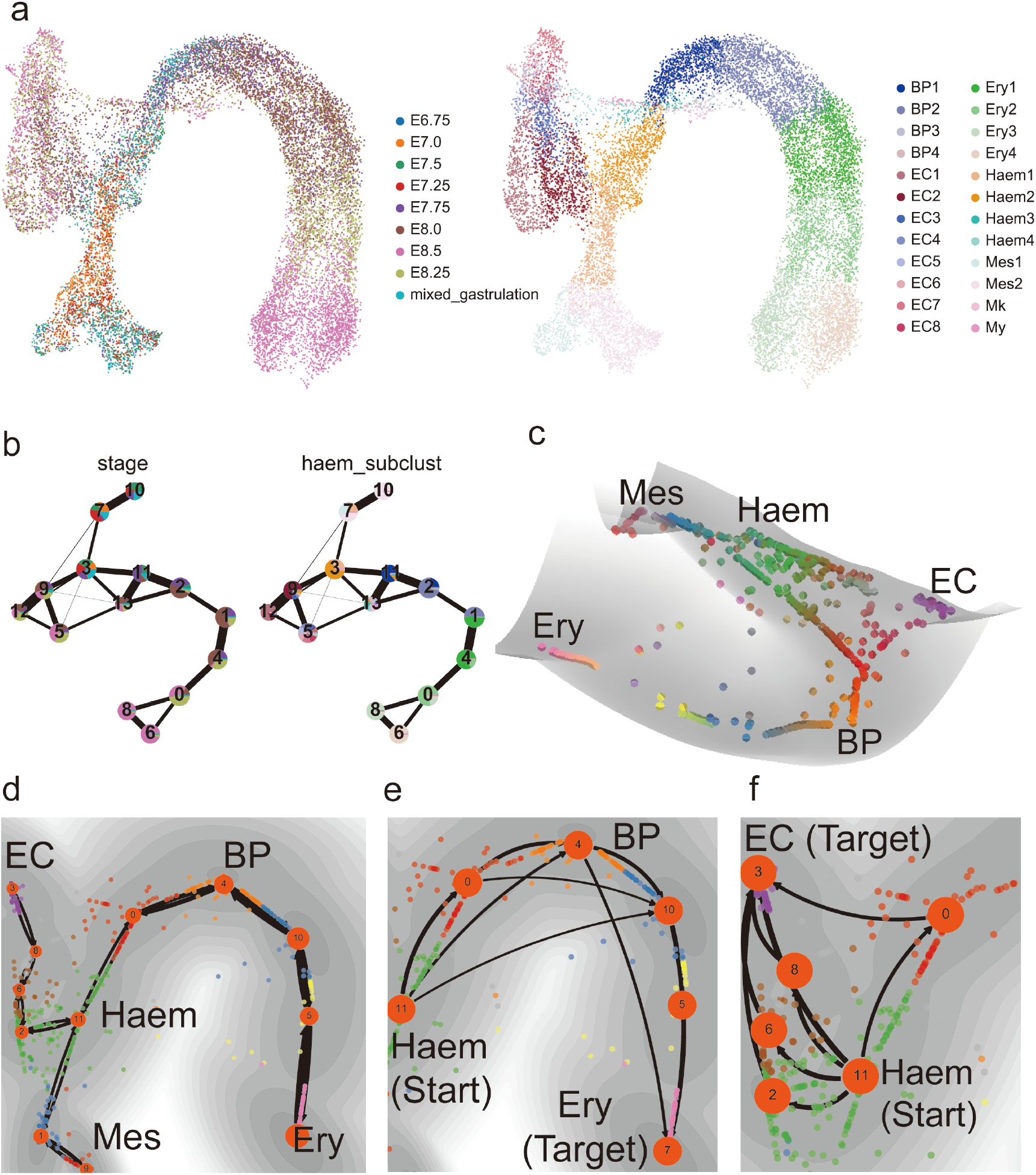
Application to a dataset on blood cell differentiation in mouse gastrulation (N=15,875 cells). (a) The UMAP plot with cells colored by experimental collection time (left) and the cell annotations in original publication (right). (b) The cell lineage inferred by PAGA, however, with the coarse-grained states colored by experimental collection time (left) and the cell annotations in the original study (right). (c) The dynamical manifold constructed by MuTrans with DECLARE pre-processing (K=1,500 micro-states), with cells colored by soft clustering membership in MuTrans attractors. (d) The global cell lineage inferred by MuTrans MPFT (most probable flow tree) algorithm. (e) Zoom-in of the dominant transition paths from Haem cells to endothelial cells. (f) Zoom-in of the dominant transition paths from Haem cells to erythrocytic cells.

### Comparison with other Methods

MuTrans is designed specifically to identify transition cells, with its associated dynamical manifold to allow easy visualization of the cell state transitions. Next we compared it with other intuitive approaches, including pseudotime ordering and cell-fate bias probability, for the detection of transition cells. We also benchmarked with seven existing methods for their capacity to unravel complex cell lineages during differentiation (**SM Section 4**).

In iPSC data, we found only MuTrans, PAGA and VarID recovered the bifurcation dynamics toward En and M states (**Figure S16**). However, the cell lineage graphs of PAGA and VarID include false-positive links that are unlikely to exist between cells collected at different time in experiments. While the projected lineage tree of StemID2 shows transition cells between precursor and mature En/M states (**Figure S16**), the reconstructed spanning tree does not reveal the overall bifurcation structure.

For myelopoiesis dataset, we found that only MuTrans and VarID constructed the bifurcations toward granulocytic or monocytic states (**Figure S17**), despite that VarID cannot distinguish the megakaryocytic and erythrocytic cells. FateID faithfully captures the differentiation paths toward monocytic states, while lacking accuracy of revealing the transitions into the granulocytic lineage (**Figure S17**).

Close inspection into the transition from precursors to mature En/M states in iPSC dataset suggests that the intuitive approaches (such as tracking the changes along pseudotime or fate bias probability) could not distinguish the transition cells from meta-stable cells as accurately and reliably as MuTrans. Both Monocle3 and DPT have a sharp increase in the pseudotime during the transitions (**Figure S18**), therefore lacking resolution in probing the transition cells linking multiple meta-stable states. Fate ID suggests a gradual change of En/M fate probability in precursor cells (**Figure S18**), not discriminating the transition cells within Pre-En and Pre-M states. Such problem was also observed when using Palantir, which depicts the entire cell-state transition as a highly continuous and gradual process (**Figure S18**).

## Discussion

Overall, MuTrans provides a unified approach to inspect cellular dynamics and to identify transition cells directly from single-cell transcriptome data across multiple scales. Central to the method is an underlying stochastic dynamic system that naturally connects attractor basins with meta-stable states, saddle points with transient states, and most probable paths with cell lineages. Instead of the widely used low-dimensional geometrical manifold approximation for the high-dimensional single-cell data, our method constructs a novel cell-fate dynamical manifold to visualize dynamics of cells development, allowing direct characterization of transition cells that move across barriers amid different meta-stable basins. Adopting the transition path theory to the multiscale dynamical system, we quantify the relative likelihoods of various transition trajectories that connect a chosen root state and the target meta-stable states. In addition, we provide a quantitative methodology to detect critical genes that drive transitions or mark meta-stable cells.

In this study a key theoretical assumption for modeling cell-state transition is a barrier-crossing picture in multi-stable dynamical systems, a concept which has been adopted previously (3, 34, 35). Indeed, the “barriers”, “saddles” and “potential landscape” underlying the actual biological process are the emergent properties of the complex interactions, such as gene expression regulation and signal transduction during a developmental process (36). The driving force that overcomes the barrier and induces the transition may arise from both the extrinsic environment and the fluctuations within the cells (37). Multi-scale reductions used by MuTrans naturally capture the transition cells, allowing inference of the corresponding transition processes.

Methods such as Palantir (31), Population Balance Analysis (PBA) (38) and Topographer (39) also treat cell-fate transition as the Markov random walk process. Unlike MuTrans, these methods only depict the dynamics at the individual cell level, lacking the capability of MuTrans to 1) resolve the intrinsic multiscale features of the system, 2) distinguish between meta-stable and transition cells, and 3) quantify the complex routes of development paths. Several other methods (2, 40) define the transition probability between clusters based on entropy difference or cell-cell transition probabilities. In comparison, the cluster-cluster scale transition probability in MuTrans is an emergent multiscale quantity derived from coarse-graining procedure, quantitatively consistent with Kramers’ reaction rate theory for over-damped Langevin dynamics (**Methods and SM**). By using such approach on transition cells, we are able to reconcile previously noted discrepancies in blood differentiation via analyzing three different datasets collected by different sequencing technologies.

Pseudotime ordering may serve as an intuitive tool to trace the progression of cell state transitions by comparing similarity of the gene expression among cells. Such approaches often adopt the deterministic point of view on cell-fate transitions, failing to distinguish between transition and meta-stable cells (**Figure 1a and S19**). In contrast, MuTrans embraces the stochastic model of cell-state transition. While cells reside and fluctuate within meta-stable states for the majority of time, it is the temporal ordering of transient transition cells, rather than meta-stable cells, reflect the actual process of cell transitions (**Figure 1c and Figure S19**).

To describe the smooth state transitions, several other methods (41, 42) adopt the soft-clustering strategy based on the soft K-means or factor decomposition for gene expression matrix. In comparison, the soft cell assignment of MuTrans is obtained from multiscale learning of cell-cluster rwTPM, which can be more robust against technical noise than using gene expression matrix directly for clustering (7). Such robustness is critical to detecting transition cells in datasets with lower sequencing depth, such as 10X data. Beyond interpreting the soft membership function as the indicator of cell locations in attractor basins, it remains an interesting problem to derive its continuous limit in the embedded over-damped Langevin dynamical systems.

To deal with the emerging large-scale scRNA-seq datasets, MuTrans introduces a pre-processing method (DECLARE) to aggregate the cells and speed up computation. The aggregation method uses the coarse-grain approach consistent with MuTrans, and it is different from other methods often used for large scRNA-seq datasets, such as down-sampling convolution (43) or kNN partition (44) that is based on the averaging or summation of cells with similar gene expression profiles. As a result, DECLARE can be naturally integrated with dynamical manifold construction and transition trajectory inference.

Admittedly, the physical picture of MuTrans cannot explain all the possible cell transition scenarios. For instance, the barrier-crossing mechanism is not sufficient to capture the oscillatory processes such as cell cycle (38). Instead of constructing cell-cell scale random walk with a pure diffusion-like kernel on transcriptome data, such non-equilibrium process might be accounted for by single-cell RNA velocity (18, 45, 46), thereafter a multi-scale reduction approach can naturally apply (47). Effective ways in root cell states detection (e.g. through entropy methods (48) or RNA velocity (46)) can also enhance the robustness of our method.

In the meantime, the back and forth stochastic transitions among meta-stable states may need to be combined with deterministic processes in order to better understand the cell-fate decision (49). The local fluctuations of microscopic cell states in gene expression can be prevalent in the dynamics, and the cell-cell scale random walk becomes a natural assumption. In theory, the stochastic transition model is consistent with the uni-direction process if the transition probabilities in one direction are dominant or when the noise amplitude of system is relatively small.

In addition to infer complex cellular dynamics induced by transition cells from single-cell transcriptome data, MuTrans along with its computational or theoretical components can be used for development of other approaches for dissecting cell-fate transitions from both data-driven and model-based perspectives.

## Methods

MuTrans performs three major tasks in order to reveal the dynamics underneath single-cell transcriptome data (**Figure 1**): 1) assigning each cell in the attractor basins of an underlining dynamical system, 2) quantifying the barrier heights across the attractor basins, and 3) identifying relative positions of the cells within each attractor. The first two tasks are executed simultaneously through the coarse-graining of multi-scale cellular random walks, an alternative approach to the traditional clustering of cells and inference of cell lineage. The third task is achieved by refining the coarse-grained dynamics via soft clustering, and serves as a critical procedure to identifying the transition cells during cell-fate conversion.

### Multi-scale analysis of the random-walk transition probability matrix (rwTPM)

We assume the underlying stochastic dynamics during cell-fate conversion be modeled by random walks among individual cells through the random-walk transition probability matrix (**rwTPM**). Dependent on the choices of either cell-level or cluster-level, the rwTPM can be constructed in different resolutions, exhibiting multi-scale property and leading the identification of transition cells from the meta-stable cells.

In describing the method, we use the indices *x,y,z* to denote individual cells and *i, j, k* to represents the clusters (or cell states) for the simplicity of notations.

#### The rwTPM in the cell-cell resolution

The rwTPM *p* of cellular stochastic transition can be directly constructed from the gene expression matrix in cell-cell resolution, with the form

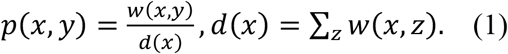

where the weight *w*(*x,y*) denotes the affinity of gene expression profile in cell *x* and *y* (**Section 2.1 in SM**). Such microscopic random walk yields an equilibrium probability distribution 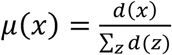, satisfying the detailed-balance condition *μ*(*x*)*p*(*x,y*) = *μ*(*y*)*p*(*y, x*). The rwTPM captures the cellular transition in the cell-cell resolution (**Figures 1d**).

#### The rwTPM in the cluster-cluster resolution

The cellular transition rwTPM can be lifted in the cluster-cluster resolution by adopting a macroscopic perspective. For example, the cell-to-cell rwTPM can be generated from certain coarse-grained dynamics, by assigning each cell in different clusters 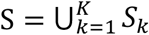, and model the transitions as the Markov Chain among clusters with the transition probability matrix 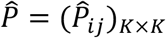. Here 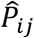 denote the probability that the cells reside in the state of cluster *S_i_* switch to the state of cluster *S_j_*. Denote 1*_S_k__*(*z*) as the indicator function of cluster *S_k_* such that 1*_S_k__*(*z*) = 1 for cell *z* ∈ *S_k_* and 1*_S_k__*(*z*) = 0 otherwise. The cluster-cluster transition based on probability matrix 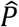 can naturally induce another rwTPM 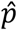 with the form

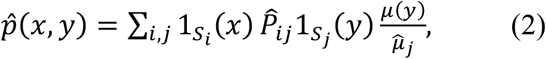

where 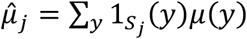 is the stationary probability distribution of cluster *S_j_*. Intuitively, the stochastic transition from cell *x* ∈ *S_i_* to *y* ∈ *S_j_* can be decomposed into a two-stage process: a cell switches cellular state from cluster *S_i_* to *S_j_* with probability 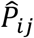, and then becomes the cell *y* in cluster *S_j_* according to its relative portion at equilibrium 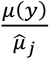. The rwTPM captures the cellular transition in the cluster-cluster resolution (**Figures 1d**).

#### The rwTPM in the cell-cluster resolution

Because some cells, for example the transition cells, may not be characterized by their locations in one basin, we introduce a membership function *ρ*(*x*) = (*ρ*_1_(*x*), *ρ*_2_(*x*),…, *ρ_K_*(*x*))*^T^* for each cell *x* to quantify its uncertainty in clustering. The element *ρ_k_*(X) represents the probability that the cell *x* belongs to cluster 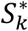 with ∑_*k*_*ρ*_*k*_(*x*) = 1. For the cell possessing mixed cluster identities, its membership function *ρ*(*x*) might have several significant positive components, suggesting its potential origin and destination during the transition process. In terms of dynamical system interpretation, the membership function captures the finite-noise effect in over-damped Langevin equation, which introduces the uncertainty of transition paths across saddle points (50), revealing that cells near saddle points and stable points may exhibit different behaviors in the state-transition dynamics.

From the coarse-grained dynamics 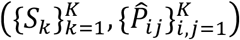 and the measurement of cell identity uncertainty *ρ_k_*(*x*) in the clusters, one can reinterpret the induced microscopic random walk 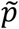 in a cell-cluster resolution as

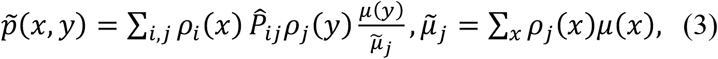

in parallel to Equation (2). Now the transition from cell *x* to *y* is realized in all the possible channels from attractor basin *S_i_* to *S_j_* with the probability *ρ_i_*(*x*)*ρ_j_*(*y*). The underlying rationale is that the transition can be decomposed in a three-stage process: First we pick up cell starting in attractor basin with membership probability, then conduct the transition with coarse-grained probability between attractor basins, and finalize the process by picking the target cell with membership probability in the target attractor basin. Now the rwTPM captures cellular transition in the cell-cluster resolution (**Figures 1d).**

#### Integrating the rwTPM at three levels

To integrate the rwTPM from different resolutions, we next optimize the rwTPM on cluster-cluster and cell-cluster level through approximating the original rwTPM in the cell-cell resolution. First, we seek an optimal coarse-grained reduction that minimizes the distance between 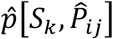 and *p* by solving an optimization problem:

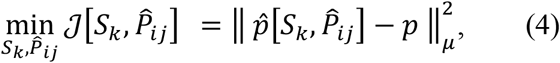

where *μ* is the stationary distribution of original cell-cell random walk *ρ*, and || ||_*μ*_ is the Hilbert-Schmidt norm (51) for transition probability matrix 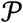, defined as 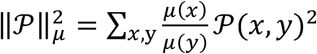. The optimization problem is solved via an iteration scheme for *S_k_* and 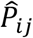 respectively (**Section 2 in SM**). The optimal coarse-grained approximation 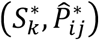 indicates the distinct clusters of cells and their mutual conversion probability. Provided with the starting state, we can infer the cell lineage from the Most Probable Path Tree (MPPT) approach or Maximum Probability Flow Tree (MPFT) approach (**Section 2 in SM**).

Next, we optimize the membership *ρ_k_*(*x*) such that the distance between the cell-cluster rwTPM 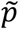 and the original *ρ* is minimized, i.e.

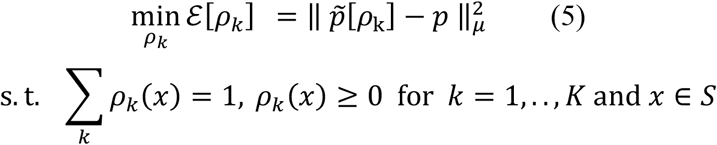

with the initial condition 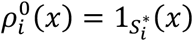, and 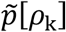 is defined from (3) by plugging in the obtained 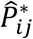. The optimization problem is solved by the quasi-Newton method (**Section 2.2 in SM**). The obtained membership function *ρ*\*(*x*) specifies the relative position of the cells within each attractor basin and is optimal in the sense that it guarantees the closest approximation of cell-cluster level rwTPM toward the cell-cell level transition dynamics.

### Transition Paths Quantification and Comparison

To quantify the cell lineages we use the transition path theory based on coarse-grained dynamics 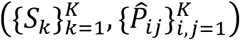 to compare the likelihood of all possible transition trajectories. Given the set of starting states *A* and the targeting state *B*, we calculate the effective current 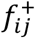 of transition paths surpassing from state *S_i_* to *S_j_* (**Section 2.4.1 in SM**), and specify the capacity of given development route *w_dr_* = (*S*_*i*_0__,*S*_*i*_1__,…,*S*_*i*_*n*__) connecting sets *A* and *B* as 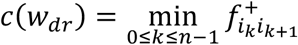. The likelihood of transition trajectory *w_dr_* is defined as the proportion of its capacity to the sum of all possible trajectory capacities. In the python package of MuTrans, we use the functions in PyEMMA (52) for the computations.

### Pre-processing by DECLARE and Scalability to Large Datasets

To reduce the computational cost for the large datasets (for instance, greater than 10K cells), we introduce a pre-processing module DECLARE (<u>d</u>ynamics-pr<u>e</u>serving <u>c</u>el<u>l</u> <u>a</u>gg<u>re</u>gation). The module first detects the hundreds/thousands of *microscopic* meta-stable states by clustering (e.g. using K-means or kNN partition) and then derive the coarse-grained transition probabilities among these *microscopic* meta-stable states. Based on such transition probabilities, we then follow the standard multiscale reduction procedure of MuTrans to find *macroscopic* meta-stable states, construct dynamical manifold, quantify the transition trajectories and highlight the transition states (**Section 2.5 in SM**).

### Transition Cells and Genes Analysis through Transcendental

Based on the soft clustering results, MuTrans performs the Transcendental (**<u>trans</u>**ition **<u>ce</u>**lls a**<u>nd</u>** r**<u>e</u>**leva**<u>nt a</u>**na**<u>l</u>**ysis) procedure to identify the transition cells from the meta-stable cells, and reveal the relevant marker genes.

For the given transition process from cluster 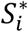 to 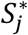 on the MPPT tree, we first selected the cells relevant to the transition, based on the membership function *ρ*\*(*x*) (**Section 2.4 in SM**). Then for each *relevant* cell *x*, we define the transition cell score (**TCS**)

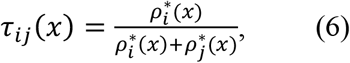

to measure the relative position of cell *x* in different clusters. Here the **TCS** *τ_ij_* takes the values near zero or one when a cell resides around the attractor in 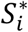 or 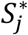 (i.e. the cells are in the meta-stable states), whereas yields the intermediate value between zero and one for the cell that possesses a hybrid or transient identity of two or more clusters. Next we arrange all the relevant cells in state 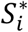 and 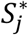 according to *τ_ij_* in descending order, and the reordered *τ_ij_* indicates a sharp transition (**Figure 1a**) or a smooth transition (**Figure 1a**) from the value one to zero. For the smooth transition, there is a group of cells whose value of *τ_ij_* decreases gradually from one to zero (**Figure 1e**). This group of cells in the transition layer are called the **transition cells** from state 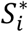 to state 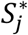, and their order reflects the details of the state-transition process. To quantify the transition steepness, we use logistic functions to model the transition and estimate the relative abundance of transition cells (**Section 2.4 in SM**). Differentially expressed genes analysis is usually applicable when the clusters are distinct and the state-transition is sharp (**Figure 1a**). However, to characterize the dynamical and hybrid gene expression profiles in transition cells, merely comparing the average gene expression in different clusters is insufficient. Here we define three kinds of genes relevant to the state transition of cells: a) the **transition-driver** (**TD**) genes that vary accordingly with the transition dynamics, b) the **intermediate-hybrid** (**IH**) genes marking the hybrid features from multiple cell states that are expressed in the intermediate transition cells, and c) the **meta-stable** (**MS**) genes that represent cells in the meta-stable states.

The expression of **TD** genes varies accordingly to the transition, revealing the driving mechanism of the cell-state conversion. To probe **TD** genes, we calculate the correlation between the gene expression values and *τ_ij_* in the ordered transition cells. The genes with larger correlation values (larger than a given threshold value) are identified as **TD** genes. The **IH** genes express eminently both in the transition cells and in the meta-stable cells from one specific cluster, reflecting the hybrid state of the transition cells, while the **MS** genes express exclusively in the meta-stable cells from certain cluster. To distinguish **IH** and **MS** genes from all the differentially expressed genes, we compare the gene expression values between the meta-stable cells and the transition cells, respectively, within each cluster. The significantly up-regulated genes in the meta-stable cells are defined as the **MS** genes, and the rest differentially expressed genes are identified as the **IH** genes that express simultaneously both in meta-stable and transition cells (**Section 2.4 in SM**).

### Constructing the cell-fate dynamical manifold

To better visualize the transition process and their connections with cell states, MuTrans introduces the dynamical manifold concept. The construction of the dynamical manifold consists of two steps: 1) locating the center positions of cell clusters (corresponding to the attractors) in low dimensional space, 2) assigning the position of each individual cells according to soft-clustering membership function.

The initial center-determination step starts with an appropriate two-dimensional representation, denoted as *x^2D^* for each cell *x* (details in **Section 2.3 in SM**). Instead of directly utilizing *x^2D^* as the cell coordinate, we calculate the center *y_k_* of each cluster 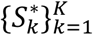 by taking the average of *x^2D^* over cells within certain range of cluster membership function 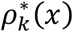. Having determined the position of attractors, we define a two-dimensional embedding *ξ*(*x*) for each cell according to the membership function *ρ*\*(*x*), such that 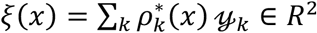. For the cell possessing mixed identities of state 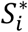 and 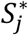, its transition coordinate then lies in a value between *y_i_* and *y_j_*. For Fokker-Planck equation of the over-damped Langevin equation, the expansion of steady-state solution near stable points (attractors) indeed yields a Gaussian-mixture distribution (53). Motivated by this, to obtain the global dynamical manifold we fit a Gaussian mixture model with a mixture weight 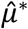 to obtain the stationary distribution of coarse-grained dynamics. The probability distribution function of the mixture model becomes

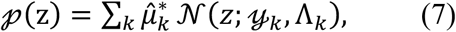

where 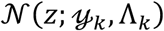 is a two-dimension Gaussian probability distribution density function with mean *y_k_* and covariance Λ*_k_*. The landscape function of dynamical manifold is then naturally takes the form in two dimensions 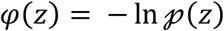. Specifically, the “energy” of individual cell *x* is calculated as *φ*(*ξ*(*x*)). The constructed landscape function captures the multi-scale stochastic dynamics of cell-fate transition, by allowing typical cells that are distinctive to certain cell states positioned in the basin around corresponding attractors, while the transition cells laid along the connecting path between attractors across the saddle point. Moreover, the relative depth of the attractor basin reflects the stationary distribution of coarse-grained dynamics, depicting the relative stability of the cell states. The flatness of the attractor basin also reveals the abundance and distribution of transition cells, indicating the sharpness of cell fate switch.

### Mathematical Analysis of MuTrans

With the assumption that the single-cell data is collected from the probability distribution *v*(*x*) with density of Boltzmann-Gibbs form, i.e., 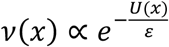, we can prove (**Section 1 in SM**) that the microscopic random walk constructed by MuTrans approximates the dynamics of over-damped Langevin Equation (OLE)

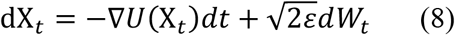

in the limiting scheme, and the coarse-graining of MuTrans 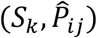 is equivalent to the model reduction of OLE by Kramers’ rate formula in the small noise regime, i.e. 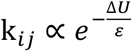 as ε → 0, where k*_ij_* is the switch rate from attractor *S_i_* to *S_j_*, and Δ*U* denotes the corresponding barrier height of transition -- the energy difference between saddle point and the departing attractor.

Therefore, if the cell transition dynamics can be well-modelled by the OLE dynamics of Equation (8), MuTrans is indeed the multi-scale model reduction of (8) via the data-driven approach. In addition, the dynamical manifold constructed by MuTrans can be viewed as the data realization of potential landscape (34) for diffusion process in biochemical modelling, which incorporates the dynamical clues about the underlying stochastic system regarding the stationary distribution and transition barrier heights.

## Data availability

All the datasets used in this paper are publicly available. The mouse cancer EMT data (Smart-Seq2) is from GSE110357, mouse myelopoiesis data (Fluidigm C1) from GSE7024, mouse hematopoietic progenitors data (Cel-Seq2) from GSE100037, human hematopoietic progenitors data (10X Chromium) from the data link in original publication (31), blood differentiation data (10X Chromium) in mouse gastrulation from https://github.com/MarioniLab/EmbryoTimecourse2018, and iPSC differentiation data (single-cell RT-qPCR) downloaded from https://www.pnas.org/highwire/filestream/29285/field_highwire_adjunct_files/1/pnas. 1621412114.sd02.xlsx. The codes and trajectories for simulation data, the processed single-cell data expression matrix, the MuTrans package and scripts to reproduce the figures and results in main text and repeat the detailed analysis in SI are also available at Github (https://github.com/cliffzhou92/MuTrans-release).

## Code availability

The Matlab implementation of MuTrans and affiliated Transcendental packages are available from GitHub (https://github.com/cliffzhou92/MuTrans-release). The Python package for MuTrans (pyMuTrans) compatible with AnnData object is also available in the repository.

## Acknowledgements

We thank Dr. Suoqin Jin and Jifan Shi for helpful discussions. We also thank the reviewers for their insightful suggestions. This project was supported by grants from the National Natural Science Foundation of China (11825102 and 11421101 to T.L.), National Institutes of Health grant U01AR073159 (Q.N.), National Science Foundation grants DMS1763272 (Q.N.) and MCB2028424 (Q.N.), and The Simons Foundation (594598 to Q.N.) of USA. T.L. is also partially supported by the Beijing Academy of Artificial Intelligence (BAAI). P.Z. also received the support from Study Abroad Program and Elite Program of Computational and Applied Mathematics for Ph.D. students of Peking University.

## Author contributions

Q.N., T.L. and P.Z. conceived the project; P.Z. and T.L. designed the algorithm and wrote the code; P.Z. and S.W. conducted the data analyses; P.Z. wrote the supplementary material; all the authors wrote and approved the manuscript. Q.N. and T.L. supervised the research.

## References

1. Svensson V, Vento-Tormo R, Teichmann SA. Exponential scaling of single-cell RNA-seq in the past decade. Nature protocols. 2018;13(4):599-.

2. Jin S, MacLean AL, Peng T, Nie Q. scEpath: energy landscape-based inference of transition probabilities and cellular trajectories from single-cell transcriptomic data. Bioinformatics. 2018;34(12):2077–86.

3. Brackston RD, Lakatos E, Stumpf MPH. Transition state characteristics during cell differentiation. PLoS Computational Biology. 2018;14(9):e1006405.

4. Moris N, Pina C, Arias AM. Transition states and cell fate decisions in epigenetic landscapes. Nature Reviews Genetics. 2016;17(11):693–703.

5. MacLean AL, Hong T, Nie Q. Exploring intermediate cell states through the lens of single cells. Current Opinion in Systems Biology. 2018;9:32–41.

6. Ohgushi M, Sasai Y. Lonely death dance of human pluripotent stem cells: ROCKing between metastable cell states. Trends in Cell Biology. 2011;21(5):274–82.

7. Haghverdi L, Buttner M, Wolf FA, Buettner F, Theis FJ. Diffusion pseudotime robustly reconstructs lineage branching. Nature Methods. 2016;13(10):845–8.

8. Sha Y, Haensel D, Gutierrez G, Du H, Dai X, Nie Q. Intermediate cell states in epithelial-to-mesenchymal transition. Phys Biol. 2019;16(2):021001.

9. Luecken MD, Theis FJ. Current best practices in single-cell RNA-seq analysis: a tutorial. Mol Syst Biol. 2019;15(6):e8746.

10. Ho YJ, Anaparthy N, Molik D, Mathew G, Aicher T, Patel A, et al. Single-cell RNA-seq analysis identifies markers of resistance to targeted BRAF inhibitors in melanoma cell populations. Genome Research. 2018;28(9):1353–63.

11. Kiselev VY, Kirschner K, Schaub MT, Andrews T, Yiu A, Chandra T, et al. SC3: consensus clustering of single-cell RNA-seq data. Nature Methods. 2017;14(5):483–6.

12. Wang B, Zhu J, Pierson E, Ramazzotti D, Batzoglou S. Visualization and analysis of single-cell RNA-seq data by kernel-based similarity learning. Nat Methods. 2017;14(4):414–6.

13. Herring CA, Banerjee A, McKinley ET, Simmons AJ, Ping J, Roland JT, et al. Unsupervised Trajectory Analysis of Single-Cell RNA-Seq and Imaging Data Reveals Alternative Tuft Cell Origins in the Gut. Cell Systems. 2018;6(1):37–51 e9.

14. Street K, Risso D, Fletcher RB, Das D, Ngai J, Yosef N, et al. Slingshot: cell lineage and pseudotime inference for single-cell transcriptomics. BMC Genomics. 2018;19(1):477.

15. Qiu X, Mao Q, Tang Y, Wang L, Chawla R, Pliner HA, et al. Reversed graph embedding resolves complex single-cell trajectories. Nat Methods. 2017;14(10):979–82.

16. Zhu L, Lei J, Klei L, Devlin B, Roeder K. Semisoft clustering of single-cell data. Proceedings of the National Academy of Sciences. 2019;116(2):466–71.

17. Zhou P, Gao X, Li X, Li L, Niu C, Ouyang Q, et al. Stochasticity Triggers Activation of the S-phase Checkpoint Pathway in Budding Yeast. Physical Review X. 2021;11(1):011004.

18. Qiu X, Zhang Y, Yang D, Hosseinzadeh S, Wang L, Yuan R, et al. Mapping vector field of single cells. Biorxiv. 2019:696724.

19. Gillespie DT. The chemical Langevin equation. The Journal of Chemical Physics. 2000;113(1):297–306.

20. Aurell E, Sneppen K. Epigenetics as a First Exit Problem. Physical Review Letters. 2002;88(4):048101.

21. Ferrell James E. Bistability, Bifurcations, and Waddington’s Epigenetic Landscape. Current Biology. 2012;22(11):R458–R66.

22. Farrell JA, Wang Y, Riesenfeld SJ, Shekhar K, Regev A, Schier AF. Single-cell reconstruction of developmental trajectories during zebrafish embryogenesis. Science. 2018;360(6392).

23. Wagner DE, Weinreb C, Collins ZM, Briggs JA, Megason SG, Klein AM. Single-cell mapping of gene expression landscapes and lineage in the zebrafish embryo. Science. 2018;360(6392):981–7.

24. Van Kampen NG. Stochastic processes in physics and chemistry: Elsevier; 1992.

25. Shi J, Li T, Chen L. Towards a critical transition theory under different temporal scales and noise strengths. Physical Review E. 2016;93(3):032137.

26. Pastushenko I, Brisebarre A, Sifrim A, Fioramonti M, Revenco T, Boumahdi S, et al. Identification of the tumour transition states occurring during EMT. Nature. 2018;556(7702):463-+.

27. Bargaje R, Trachana K, Shelton MN, McGinnis CS, Zhou JX, Chadick C, et al. Cell population structure prior to bifurcation predicts efficiency of directed differentiation in human induced pluripotent cells. Proceedings of the National Academy of Sciences. 2017;114(9):2271–6.

28. Jia C, Zhang MQ, Qian H. Emergent Levy behavior in single-cell stochastic gene expression. Phys Rev E. 2017;96(4-1):040402.

29. Olsson A, Venkatasubramanian M, Chaudhri VK, Aronow BJ, Salomonis N, Singh H, et al. Single-cell analysis of mixed-lineage states leading to a binary cell fate choice. Nature. 2016;537(7622):698–702.

30. Herman JS, Sagar, Grun D. FateID infers cell fate bias in multipotent progenitors from single-cell RNA-seq data. Nat Methods. 2018;15(5):379–86.

31. Setty M, Kiseliovas V, Levine J, Gayoso A, Mazutis L, Pe’er D. Characterization of cell fate probabilities in single-cell data with Palantir. Nat Biotechnol. 2019;37(4):451–60.

32. Wolf FA, Hamey FK, Plass M, Solana J, Dahlin JS, Gottgens B, et al. PAGA: graph abstraction reconciles clustering with trajectory inference through a topology preserving map of single cells. Genome Biol. 2019;20(1):59.

33. Pijuan-Sala B, Griffiths JA, Guibentif C, Hiscock TW, Jawaid W, Calero-Nieto FJ, et al. A single-cell molecular map of mouse gastrulation and early organogenesis. Nature. 2019;566(7745):490–5.

34. Wang J, Zhang K, Xu L, Wang E. Quantifying the Waddington landscape and biological paths for development and differentiation. P Natl Acad Sci USA. 2011;108(20):8257–62.

35. Zhou P, Li T. Construction of the landscape for multi-stable systems: Potential landscape, quasi-potential, A-type integral and beyond. The Journal of Chemical Physics. 2016;144(9):094109.

36. Huang S, Li F, Zhou JX, Qian H. Processes on the emergent landscapes of biochemical reaction networks and heterogeneous cell population dynamics: differentiation in living matters. J R Soc Interface. 2017;14(130).

37. Elowitz MB, Levine AJ, Siggia ED, Swain PS. Stochastic gene expression in a single cell. Science. 2002;297(5584):1183–6.

38. Weinreb C, Wolock S, Tusi BK, Socolovsky M, Klein AM. Fundamental limits on dynamic inference from single-cell snapshots. Proc Natl Acad Sci U S A. 2018;115(10):E2467–E76.

39. Zhang J, Nie Q, Zhou T. Revealing Dynamic Mechanisms of Cell Fate Decisions From Single-Cell Transcriptomic Data. Front Genet. 2019;10:1280.

40. Grun D. Revealing dynamics of gene expression variability in cell state space. Nat Methods. 2020;17(1):45–9.

41. Zheng X, Jin S, Nie Q, Zou X. scRCMF: Identification of cell subpopulations and transition states from single cell transcriptomes. IEEE Trans Biomed Eng. 2019.

42. Korsunsky I, Millard N, Fan J, Slowikowski K, Zhang F, Wei K, et al. Fast, sensitive and accurate integration of single-cell data with Harmony. Nat Methods. 2019;16(12):1289–96.

43. Iacono G, Mereu E, Guillaumet-Adkins A, Corominas R, Cusco I, Rodriguez-Esteban G, et al. bigSCale: an analytical framework for big-scale single-cell data. Genome Res. 2018;28(6):878–90.

44. Baran Y, Bercovich A, Sebe-Pedros A, Lubling Y, Giladi A, Chomsky E, et al. MetaCell: analysis of single-cell RNA-seq data using K-nn graph partitions. Genome Biol. 2019;20(1):206.

45. La Manno G, Soldatov R, Zeisel A, Braun E, Hochgerner H, Petukhov V, et al. RNA velocity of single cells. Nature. 2018;560(7719):494–8.

46. Bergen V, Lange M, Peidli S, Wolf FA, Theis FJ. Generalizing RNA velocity to transient cell states through dynamical modeling. Nat Biotechnol. 2020;38(12):1408–14.

47. Li T, Shi J, Wu Y, Zhou P. On the Mathematics of RNA Velocity I: Theoretical Analysis. bioRxiv. 2020.

48. Shi J, Teschendorff AE, Chen W, Chen L, Li T. Quantifying Waddington’s epigenetic landscape: a comparison of single-cell potency measures. Brief Bioinform. 2018.

49. Guillemin A, Roesch E, Stumpf MPH. Uncertainty in cell fate decision making: Lessons from potential landscapes of bifurcation systems. bioRxiv. 2021:2021.01.03.425143.

50. Pinski F, Stuart A. Transition paths in molecules at finite temperature. The Journal of Chemical Physics. 2010;132(18):184104.

51. E W, Li T, Vanden-Eijnden E. Optimal partition and effective dynamics of complex networks. Proceedings of the National Academy of Sciences. 2008;105(23):7907–12.

52. Scherer MK, Trendelkamp-Schroer B, Paul F, Pérez-Hernández G, Hoffmann M, Plattner N, et al. PyEMMA 2: A Software Package for Estimation, Validation, and Analysis of Markov Models. Journal of Chemical Theory and Computation. 2015;11(11):5525–42.

53. Pearce P, Woodhouse FG, Forrow A, Kelly A, Kusumaatmaja H, Dunkel J. Learning dynamical information from static protein and sequencing data. Nature Communications. 2019;10(1):5368.

